# Draft genome sequences of five Proteobacteria isolated from Lechuguilla Cave, New Mexico, USA: Insights into taxonomy and quorum-sensing

**DOI:** 10.1101/742460

**Authors:** Han Ming Gan, Peter C. Wengert, Hazel A. Barton, André O. Hudson, Michael A. Savka

## Abstract

Genomic resources remain scarce for bacteria isolated from oligotrophic caves. We sequenced the genomes of five Proteobacteria isolated from Lechuguilla Cave in New Mexico, USA. Genome-based phylogeny indicates that each strain belongs to a distinct genus. Two *Rhizobiaceae* isolates possess the genomic potential for the biosynthesis of acyl-homoserine lactone.

## Main Text

Adequate genomic resources are crucial for the understanding of adaptation for culturable microbes in nutrient-limited cave environments (1). To date, bacterial genomes isolated from caves are poorly represented in the public databases (2-5). Here, we report the genomes of five Lechuguilla cave isolates and used genome-based phylogeny to refine their taxonomic assignment. We also identified genomic potential for the biosynthesis of cell-cell communication signal in two *Rhizobiaeae* isolates and confirmed the predicted phenotype using an *Agrobacterium tumefaciens* reporter assay (6).

Initial isolation of the bacterial strains from remote sample sites in Lechuguilla Cave was previously described by Bhullar *et al* (7). Strains were grown in half strength tryptic soy broth with shaking at 30°C for 2 days. DNA extraction used the GenElute bacterial genomic DNA kit (MilliporeSigma, St Louis, MO). Sequencing library was generated using the tagmentation-based Illumina Nextera XT DNA sample prep kit (Illumina, San Diego, CA) and sequenced on an Illumina MiSeq (run configuration of 2 × 150 bp). The paired-end reads were adapter-trimmed and assembled with Trimmomatic v0.36 (8) and Unicycler v0.4.7 (9), respectively, using the default setting.

The genome of LC387 was assembled into a single contig (Table 1). Based on BLASTN alignment (10), its full-length 16S rRNA sequence is 100% identical to that of *Afipia massiliensis* CCUG 45153 ^T^ (RefSeq Accession: NR_025646.1). For the generation of genome-based phylogeny using GToTree v1.2.1 (11), genome assemblies of bacterial type strains exhibiting high 16S rRNA gene sequence similarity to the cave isolates were downloaded and included in the pipeline. Based on phylogenomic clustering, confident taxonomic assignment to the genus level was obtained for LC34, LC103 and LC458 (Figure 1A). The basal placement of LC148 in the *Neorhizobium* clade suggests that it is a divergent member within the genus or a member of an undescribed genus. Therefore, strain LC148 was classified as *Rhizobacteriaceae* sp. pending future taxonomic investigation.

**Table 1.**
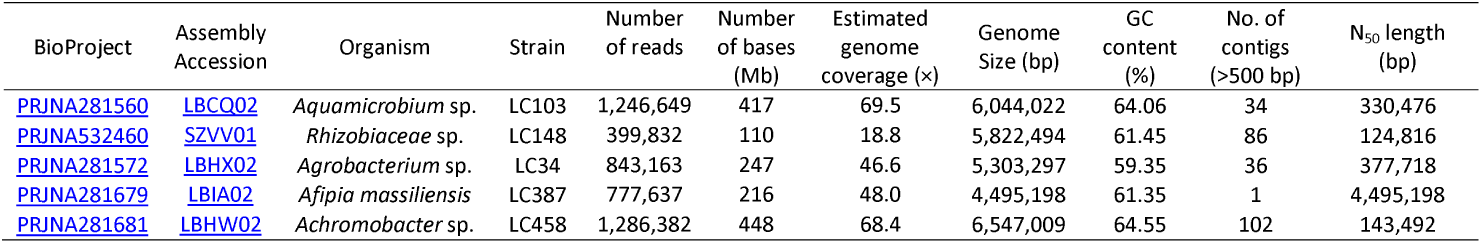
Genome assembly metrics and data availability.

| BioProject | Assembly<br>Accession | Organism | Strain | Number<br>of reads | Number<br>of bases<br>(Mb) | Estimated<br>genome<br>coverage (x) | Genome<br>Size (bp) | GC<br>content<br>(%) | No. of<br>contigs<br>(>500 bp) | N<sub>50</sub> length<br>(bp) |
| --- | --- | --- | --- | --- | --- | --- | --- | --- | --- | --- |
| PRJNA281560 | LBCQ02 | *Aquamicrobium* sp. | LC103 | 1,246,649 | 417 | 69.5 | 6,044,022 | 64.06 | 34 | 330,476 |
| PRJNA532460 | SZVV01 | *Rhizobiaceae* sp. | LC148 | 399,832 | 110 | 18.8 | 5,822,494 | 61.45 | 86 | 124,816 |
| PRJNA281572 | LBHX02 | *Agrobacterium* sp. | LC34 | 843,163 | 247 | 46.6 | 5,303,297 | 59.35 | 36 | 377,718 |
| PRJNA281679 | LBIA02 | *Afipia massiliensis* | LC387 | 777,637 | 216 | 48.0 | 4,495,198 | 61.35 | 1 | 4,495,198 |
| PRJNA281681 | LBHW02 | *Achromobacter* sp. | LC458 | 1,286,382 | 448 | 68.4 | 6,547,009 | 64.55 | 102 | 143,492 |

**Figure 1.**
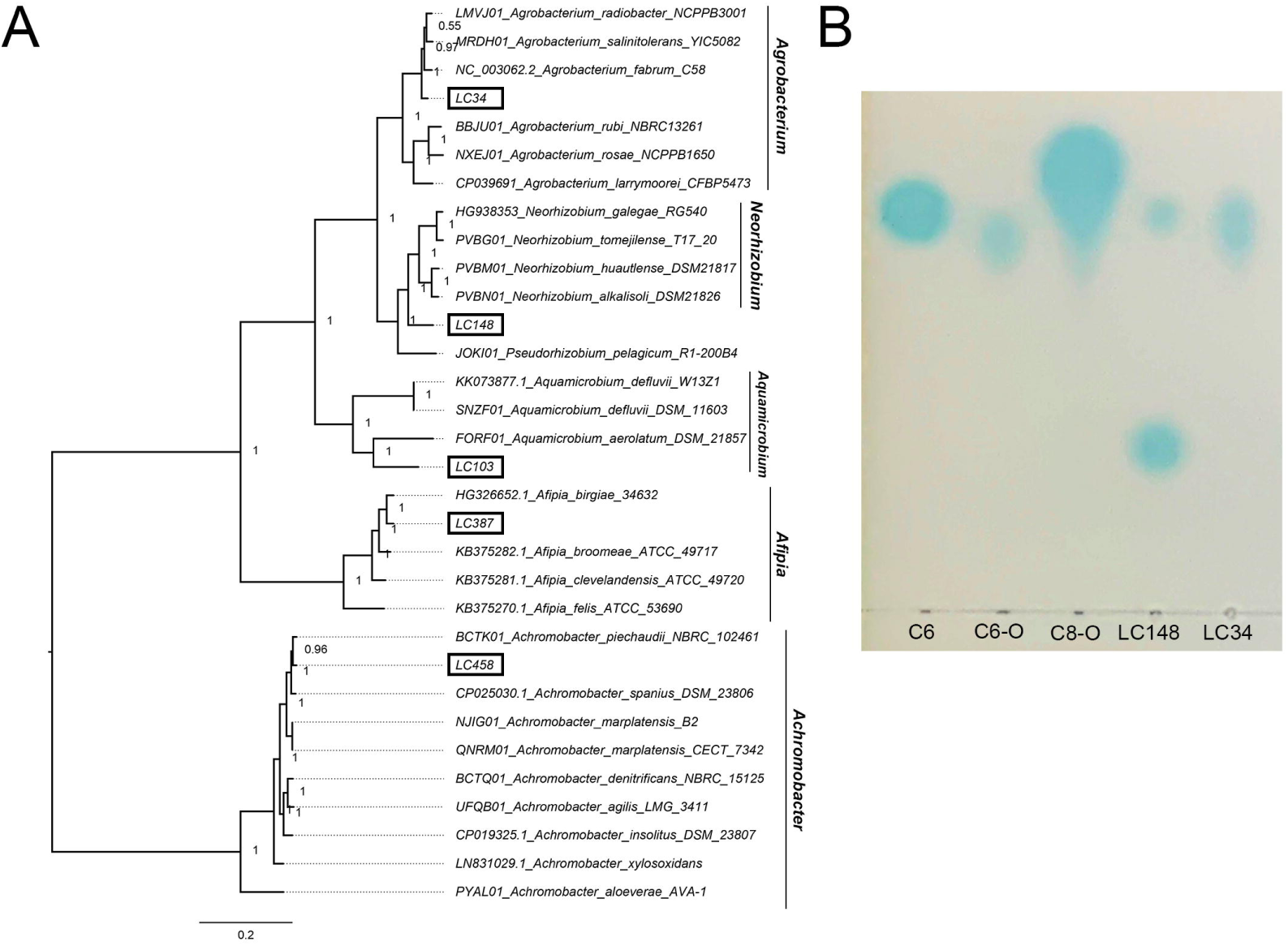
(A) Maximum likelihood tree based on the concatenated alignment of 119 conserved proteobacteria single-copy gene sets generated using the default setting of GToTree v1.2.1. Branch lengths indicate the number of substitutions per site while node labels are Shimodaira-Hasegawa (SH) local support values computed in FastTree2 (20). (B) Separation of AHL molecules from the LC34 and LC148 extracts by C_18_ reversed-phase thin-layer plate developed with methanol/water (60:40, vol/vol). Standards include: C6, N-hexanoyl-L-homoserine lactone; C6-O, N-(3-oxohexanoyl)-L-homoserine lactone; C8-O, N-(3-oxo-octanoyl)-L-homoserine lactone.

We used a previously described Hidden Markov Model (HMM) approach (12) to identify the genomic potential for biosynthesis of acyl-homoserine lactone (AHL) molecules involved in the regulation of gene expression in response to cell density (13). AHL synthase homologs (LuxI) were identified in strains LC34 (GenBank Protein ID: TKT57503.1) and LC148 (GenBank Protein IDs: TKT46115.1 and TKT66962.1). The genes coding for these proteins were similarly classified as *luxI* homologs by AntiSMASH 4 and the NCBI prokaryotic genome annotation pipeline (14, 15). To confirm the predicted phenotype, LC34 and LC148 were grown in YM media and ethyl acetate extracts from the cultures were tested for the presence of AHL signals using the AHL-dependent biosensor *Agrobacterium tumefaciens* NTL4(pZLR4) (16). LC34 produced one type of AHL signal with retardation factor (RF) that is similar to 3-oxo-C8. LC148 produced two distinct AHLs, one with RF similar to C6 and the other with RF smaller than the included AHL standards (Figure 1B). Future work investigating the role of quorum sensing in these cave isolates through transposon mutagenesis (17), targeted gene deletion (18) or transcriptome sequencing (19) will be informative in understanding bacteria from cave environments.

## Data availability

The raw Illumina paired-end reads and genome assemblies have been deposited in GenBank under the BioProject numbers as listed in Table 1. Bacterial strains can be requested from Professor Michael A. Savka, Rochester Institute of Technology, New York, USA.

## Acknowledgements

The authors acknowledge the Thomas H. Gosnell School of Life Sciences (GSoLS) and the College of Science (COS) at the Rochester Institute of Technology (RIT) for ongoing support. PCW was supported by a 2019 COS Summer Undergraduate Research Fellowship from RIT.

